# OVEREXPRESSION OF TEMPRANILLO-LIKE PROTEINS PROMOTES ENDORMANCY RELEASE IN POPLAR

**DOI:** 10.1101/2025.01.11.632540

**Authors:** D. Gómez-Soto, P. M. Triozzi, D. Conde, C. del Barrio, I. Allona, M. Perales

## Abstract

Trees in temperate and boreal latitudes synchronize their growth-dormancy cycles with seasonal environmental variations to ensure their survival over the years. Dormancy control is crucial during winter when plants cease growth and establish buds to protect their apical meristems from cold temperatures. To overcome endormancy, initiate bud break, and restore growth, plants must be exposed to a specific duration of chilling, referred to as the chilling requirement, which is species- and ecotype-dependent. In this work, we study the novel roles of two *TEMPRANILLO*-like genes (*TEML1* and *TEML2*) in the annual cycle of poplar. We demonstrated that *Populus TEML* genes are regulated by photoperiod, cold temperatures and the circadian clock, and they play a role in the control of endodormancy. Notably, their function diverges from the role of its Arabidopsis ortholog *AtTEM*, which regulates *FLOWERING LOCUS T (FT)* transcription and the photoperiodic flowering transcription. Transcriptomic analysis of endodormant buds during winter revealed that the activation of *TEML1* and *TEML2* promotes endodormancy release by modulating the expression of endodormancy regulators and growth-promoting genes.

## INTRODUCTION

Perennial plants in boreal and temperate regions interpret environmental variations to anticipate the seasons, developing annual cycles of growth and dormancy. To protect meristems from winter, deciduous trees initiate autumnal growth cessation, which culminates in the formation of winter buds and the establishment of endodormancy (Rohde et al., 2002; Cooke et al., 2012). Several processes occur to reach winter dormancy, cease cell division (Karlberg et al., 2010), downregulation of growth promoting pathways (Ruttink et al., 2007), increase ABA levels (Karlberg et al., 2010; Ruttink et al., 2007), and close the plasmodesmata (Tylewicz et al., 2018). Once the endodormancy is established, endodormant buds are insensitive to growth-promoting environmental signals and unable to break the dormancy. In fact, dormancy release is a gradual process that occurs after a prolonged exposure to low temperatures (Brunner et al., 2014; Espinosa-Ruiz et al., 2004; Hannerz et al., 2003; Saure, 1985). The timing of cold required to overcome dormancy is referred to as chilling requirement, and it differs among species and ecotypes. Once chilling requirement is fulfilled, buds re-enter in ecodormancy again and regain the ability to restore growth under optimal environmental conditions (Rohde & Bhalerao, 2007).

Chilling requirements are critical for spring growth and flowering and are seriously threatened by climate change (Luedeling et al., 2011). The internal mechanism that measures the amount of chilling hours required to release dormancy in trees remains unknown (Yang et al., 2021).

Poplar is a representative plant model species used to study growth-dormancy cycles in deciduous trees. Photoperiod and temperature, which vary depending on the geographical areas, are the main environmental factors that control the growth-dormancy transition (Maurya & Bhalerao, 2017). Under long day (LD) conditions, poplar vegetative growth, which consists of internode elongation and shoot organogenesis, persists consistently until the photoperiod shortens below a critical day length (CDL)(Weiser, 1970). This reduction in hours of light triggers growth cessation and bud set, and it is specific to the tree’s ecotype and geographical origin (Rohde et al., 2010). The regulatory mechanism by which poplar plants measure photoperiod is homologous to the CONSTANS (CO)/FLOWERING LOCUS T (FT) regulatory module that controls flowering time in response to daylength variation in Arabidopsis (Bohlenius et al., 2006). Specifically, a whole-genome duplication of FT locus in poplar has generated two functionally diverged paralogs, FLOWERING LOCUS T1 (FT1) and FLOWERING LOCUS T2 (FT2), with FT2 responsible of vegetative growth (Hsu et al., 2011). Similar to the external coincidence model for photoperiodic control of Arabidopsis flowering, the precise alignment of daylength with the circadian rhythms quantitively activates *FT2* and promotes poplar vegetative growth (Alique et al., 2024). *FT2* diurnal activation is regulated by the circadian clock activator GIGANTEA (GI) and the repressive factors CYCLING DOF FACTOR 2 (CDF2), LATE ELONGATED HYPOCOTYL 2 (LHY2), and TIMING OF CAB EXPRESSION 1 (TOC1) (Alique et al., 2024; Ding et al., 2018). Quantitative genetic population studies in *Populus tremula* show that a single genomic region containing the *FT2* gene accounts for 65% of the observed genetic variation in the timing of growth cessation and bud set (Wang et al., 2018). Genome editing knock-out studies demonstrate that *FT2* is essential to sustain summer shoot growth (André et al., 2022; Gómez-Soto et al., 2022). All these studies indicate that FT2 represents the main photoperiodic trigger for growth cessation in poplar.

In Arabidopsis, the circadian clock regulated genes TEMPRANILLO 1 (TEM1) and TEMPRANILLO 2 (TEM2) have been shown to directly repress *FT* and flowering (Castillejo & Pelaz, 2008; Hu et al., 2021). TEM1 and TEM2 are transcription factors belonging to the RELATED TO ABSCISIC ACID INSENSITIVE3 (ABI3)/VIVIPAROUS (VP1) (RAV) family (Castillejo & Pelaz, 2008). The Arabidopsis RAV family comprises six members: RAV1, RAV1-like, RAV2 (renamed as TEM1), RAV2-like (renamed as TEM2), RAV3 and RAV3-like. The main characteristic of RAV genes is the presence of two DNA binding domains, B3 and an AP2, which recognize different DNA consensus sequences (Matías-Hernández et al., 2014). Most members of the B3 super-family show a repressive activity as due to the presence of a B3 repression domain (BRD), whose core sequence is also found in four members of RAV family: TEM1, TEM2, RAV1 and RAV1-like (Matías-Hernández et al., 2014). Their role as transcriptional repressors of *FT* has been studied in different plant species. The overexpression of *TEM* results in an extremely delayed transition to flowering in Arabidopsis, accompanied by the repression of *FT* transcription in leaves. Loquat EjRAV1 and EjRAV2 can bind *EjFT1* and *EjFT2* promoters and when overexpressed in Arabidopsis delays flower initiation (Peng et al., 2021). Overexpression of *Malus domestica MdTEM1* and *MdTEM2* orthologs delays the flowering of *Fragaria vesca* plants and causes repression of the *FvFT1* gene (Dejahang et al., 2023). In addition, other functions have been described in RAV family members in perennial species. Pear PbRAV6 is also involved in the response to drought stress (Liu et al., 2021). In Castanea, CsRAV1 is associated with the formation of sylleptic branches in hybrid poplar (Moreno-Cortés et al., 2012).

In this study, we investigated the roles of TEML1 and TEML2 as transitional regulators of the annual growth-dormancy cycles in poplar. The daily expression of *TEML1* and *TEML2* is fine-tuned by the circadian clock, and both are induced by short photoperiod and low temperature. Unexpectedly, phenological assays and transcriptomic analyses under short days or low temperature using overexpressing lines for poplar TEML1 and TEML2 revealed that these transcription factors are involved in the regulation of endodormancy in poplar, suggesting a divergent function compared with their orthologs in Arabidopsis. Finally, we discussed and proposed a mechanism where the chilling-dependent induction of TEML genes promotes endodormancy release in poplar.

## MATERIAL AND METHODS

### Growth conditions and generation of poplar overexpressing lines

The hybrid poplar *Populus tremula × alba* INRA clone 717 1B4 was used as the experimental model. Poplar plantlets were grown *in vitro* in Murashige and Skoog medium 1B (pH 5.7) supplemented with 2% sucrose and with indole acetic acid (0.5 mg/L) and indole butyric acid (0,5 mg/L) containing 0.7% (w/v) plant agar under long days (LD) 16h light/ 8h dark and 22°C conditions.

The *TEML1 and TEML2* coding region sequences (CDS) were amplified from hybrid poplar genomic DNA (gDNA) using gene-specific primers containing attB sites (Supplementary Table 1) for Gateway cloning system. For PCR, Phusion DNA Polymerase (Thermo Fisher Scientific, Massachusetts, USA) was used, and the PCR products were purified and cloned into pDONR207 (Life Technologies, Carlsbad, CA, USA). Insertions in the resulting entry clones were verified by sequencing. The Gateway cassettes carrying *TEML1* and *TEML2* CDS were then cloned into the destination binary vector pGWB15 (Nakagawa *et al.,* 2007). These constructs were transformed into *Agrobacterium tumefaciens* strain GV3101/pMP90 (Konczl & Schell, 1986). Next, hybrid poplar was transformed via an *Agrobacterium*-mediated protocol described previously (Gallardo *et al.,* 1999). Selection was conducted in kanamycin-containing medium and once plantlets were regenerated, the gene expression levels of *TEML1* and *TEML2* were analyzed by quantitative reverse transcription-polymerase chain reaction (qRT-PCR) in the independent transformed individuals (lines) and in wild type individual. For the experiments conducted in this study, we selected TEML1ox3, TEML1ox4, TEML2ox6 and TEML2ox11.

### Phenological assays in poplar overexpressing lines

For the phenological assays, *in vitro* cultivated poplars of wild type (WT), TEML1ox3, TEML1ox4, TEML2ox6 and TEML2ox11 were transferred to blond peat pots (pH 4.5). Plants were grown under LD and 22 °C for 3 weeks. Plants were fertilized once every 2 weeks with a solution of 1 g l−1 Peters Professional 20-20-20 in LD conditions and 20-10-20 in short day (SD, 8h light / 16h dark) conditions (Comercial Química Massó, Barcelona, Spain). All the experiments described in this study had their own WT poplar plants as controls, were conducted independently, and were repeated twice.

Growth cessation and bud set was induced by exposing plants to SD conditions (8h light/ 16h dark) and 22°C during 10 weeks. Bud set progression was graded by scoring from stage 3 (fully growing apex) to stage 0 (fully formed apical bud) according to Rohde *et al.,* 2010. Winter conditions were emulated by treating plants under SD and 4°C for 4 or 6 weeks to fulfil the chilling requirement. Finally, plants were transferred back to LD and 22 °C to monitor bud break. The bud break was scored according to the six developmental stages (from 0 to 5) according to Johansson *et al.,* 2014. For the endodormancy evaluation, plants were exposed to SD and 4°C for 3 weeks. After that, plants were transferred to LD and 22°C conditions. Phenological assays for assessing bud break were repeated twice with similar results.

### RNA extraction, cDNA synthesis and qRT-PCR analysis

RNA extraction was performed by adding to frozen buds CTAB extraction buffer with 2% β-mercaptoethanol at 65°C for 5-8 minutes, following by 3 washes with chloroform. After that, 7.5 M LiCl_2_:50mM EDTA was added, and samples were precipitated overnight at 4°C. Next day, RNA was collected by centrifuging at 10.000 RPM for 20 minutes, the supernatant was discarded, and the pellet was resuspended using NR buffer. RNA was then purified according to the manufacturer’s instructions of NZY RNA purification kit (NZYTech, Lisboa, Portugal). RNA extraction protocol is explained in detail in Gómez-Soto *et al.,* 2021. A total of 100 ng RNA was retrotranscribed to cDNA using the Maxima First Strand cDNA Synthesis kit from Thermo Scientific (Thermo Fisher Scientific, Massachusetts, USA). The qRT-PCR procedure was conducted following previously established protocols (Ramos-Sánchez *et al.,* 2019). The gene-specific primers used in these studied are listed in the supplementary Table 1.

### Gene expression studies

For temperature and photoperiod dependency studies, *in vitro* poplar WT plants exposed to SD with a declining temperature gradient: 22°C (permissive temperature), 15°C (mild cold) and 4°C (chilling temperatures), over a duration of 1 week to monitor gene responses. Each sample consisted of two leaves from different plants, collected at 3-hour intervals. To evaluate *FT2*, *GA2ox1* and *GA3ox2* expressions, in vitro poplar leaves were collected under LD conditions every 3 hours. For the evaluation of the effect of constant light, dark hours were removed, and leaves from in vitro poplar were collected at zt18 over two days. Two biological replicates were included in the analysis.

### Analysis of comparative RNAseq libraries of wild-type, TEML1ox and TEML2ox apical buds

Apical buds of plants exposed to 10 weeks of SD and 22°C conditions and 2 weeks of SD and 4°C were collected at ZT7. A total of 100 ng of extracted RNA was used to generate the RNA-seq libraries. Libraries were generated from cDNA samples using the TruSeq Stranded mRNA Library Prep Kit (Nat Rev Genet. 2011 Sep 7;12(10):671-82) and they were sequenced on the Illumina platform. Reads were then mapped to the reference genome of *Populus tremula x alba* HAP2 v5.1 with HISAT2. Expression profiles were represented as read count data. The differential analysis of sequence read count data was conducted using the EdgeR package in R, with a significance threshold set at p value p<0.05 to identify the lists of differentially expressed genes (DEG). GO enrichment analysis was performed using gprofiler tool (https://biit.cs.ut.ee/gprofiler/gost). We created GO term-Gene networks using the igraph package in R and clustered them using the Louvain Method, where genes and GO terms with shared biological roles form connected communities.

The apical bud samples used from mid-winter to spring transcriptional profiling were described earlier (Conde *et al.,* 2019) and available at https://phytozome.jgi.doe.gov/pz/portal.html.

### In silico *FT2* promoter analysis

Promoter sequences of *PtaFT2* and *PtreFT2* (*PtXaAlbH.10G142200 and PtXaTreH.10G148700*) were obtained from Phytozome (https://phytozome-next.jgi.doe.gov/). A 2kb region upstream of the 5’UTR, along with the 5’UTR itself, was included in the promoter analysis performed using the web database Plant Pan 4.0 (https://plantpan.itps.ncku.edu.tw/plantpan4/index.html; Chow et al., 2024).

## RESULTS AND DISCUSSION

### Seasonal cues and the circadian clock influence the expression of *TEML1* and *TEML2*

To investigate whether the expression of *TEML1* and *TEML2* aligns with the regulation of the seasonal growth-dormancy transition, we examined how photoperiod and temperature, along with the circadian clock, impact the gene expression of *TEML1* and *TEML2*. We found that changes in photoperiod altered the expression patterns of both *TEML* genes in poplar in similar manner. Under long days (LD), the peak expression of *TEML* occurs around midnight, specifically at ZT20 in wild type (WT) plants (**Fig. 1A and 1D**). In contrast, under short days (SD), the expression peak shifted to ZT8-9, aligning with the day-night transition (**Fig. 1B and 1E**). Analysis of gene expression in WT and *lhy1/lhy2* mutant background plants revealed notable differences. In *lhy1/lhy2* plants grown under LD, the peak expression of *TEML1* and *TEML2* increased 3-fold and 4-fold, respectively, compared with the WT (**Fig. 1A and 1D**). Conversely, under SD conditions, the expression of both *TEML* genes was significantly repressed in *lhy1/lhy2* plants, particularly during their peak expression time (**Fig. 1B and 1E**). These findings indicate that the circadian genes *LHY1* and *LHY2* modulate *TEML* gene expression in opposing manners. Specifically, they repress expression under LD conditions while promoting expression under SD conditions. This suggests that the circadian clock plays a role in the regulation of *TEML* genes. The evaluation of the 48h expression pattern under constant light (LL) indicated the circadian regulation of *TEML1* and *TEML2.* Both genes continued to show oscillation under LL, however with a period of approximately 28h and a smaller amplitude (**Supporting Information Fig. S1**). Together, these findings imply that the daily expression pattern of *TEML* genes is influenced by both photoperiod and the circadian clock.

**Figure 1.**
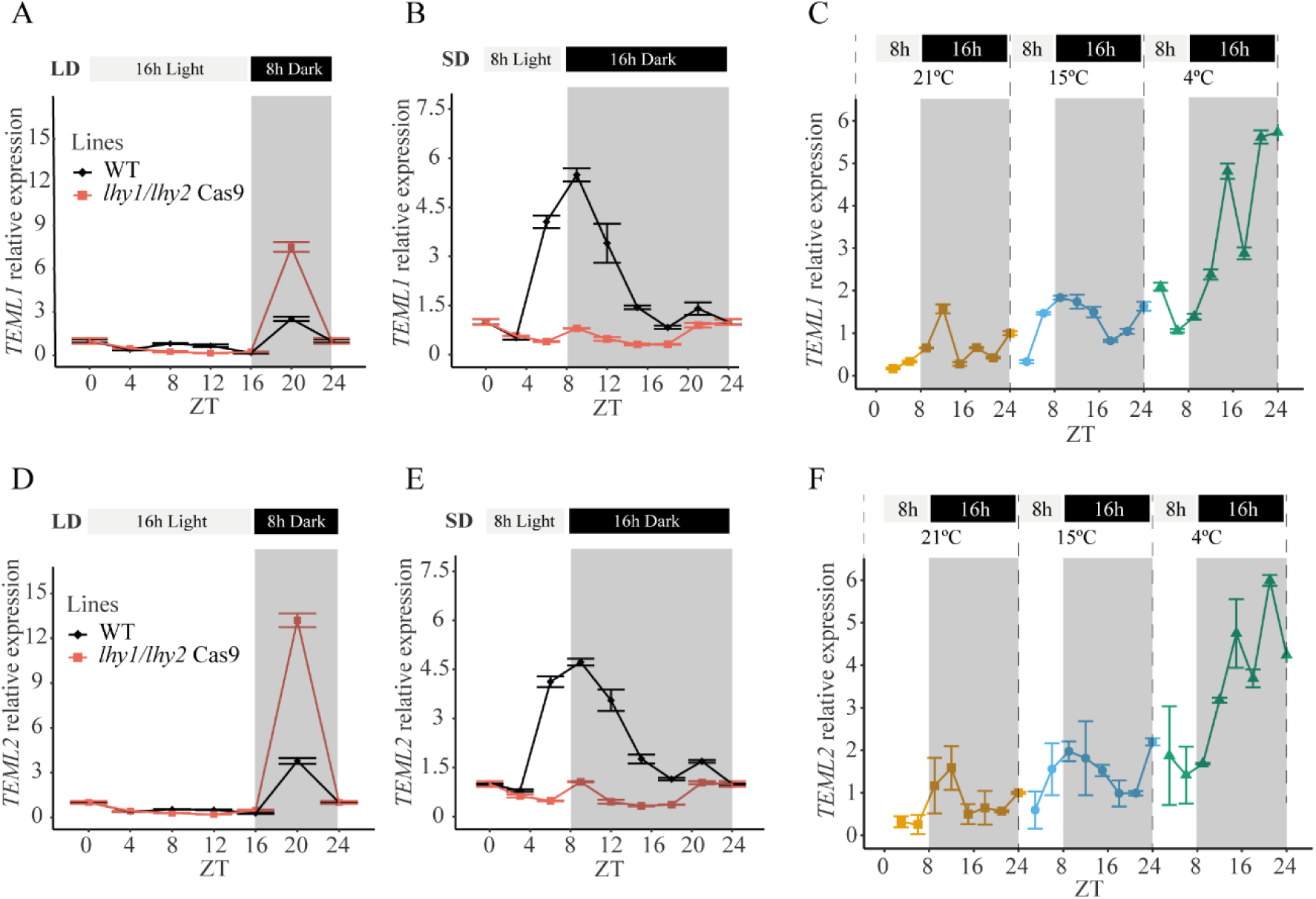
Photoperiodic and temperature regulation of *TEML1* and *TEML2*. qRT-PCR analysis of the relative expression pattern of *TEML1* (A-C) and *TEML2* (D-F) in poplar leaves. *TEML1* and *TEML2* were exposed to LD conditions (A, D), SD conditions (B, E) and a decreasing temperature gradient: 21°C, 15°C and 4°C (C, F). The top panels indicate photoperiod in which light grey boxes mean hours of light and black boxes mean hours of dark, which is also indicated with a dark background inside the plot. Samples (A, B, D, E) were collected from poplar plants grown *in vitro* every 3-4 hours for 24h. UBQ7 was used as housekeeping gene. In the case of the temperature treatment, leaves were collected after 1 week of SD and 21°C (yellow line), 15°C (blue line) or 4°C (green line). Plotted values and error bars are fold-change means + s.d. of two biological replicates.

Finally, to investigate whether low temperature regulates *TEML1* and *TEML2*, we measured their expression over 24 hours under mild cold (15°C) and chilling (4°C) conditions in SD, simulating autumnal and winter conditions, respectively. We found that the expression of both genes increased in response to cold, particularly under chilling temperatures which resulted in greater accumulation during nighttime (**Fig. 1C and 1F**). Collectively, our results show that the poplar genes *TEML1* and *TEML2* are highly sensitive to autumnal-winter cues and the circadian clock, with their daily gene expression levels induced by short days and cold temperatures.

### *TEML1* and *TEML2* overexpression do not repress *FT2* and do not affect poplar seasonal growth cessation

To investigate the role of *TEML1* and *TEML2* during the annual growth-dormancy cycle, we generated poplar *TEML1* and *TEML2* overexpressing lines and conducted phenological assays under controlled condition in growth chambers using two different overexpressing lines for each gene: TEML1ox3, TEML1ox4, TEML2ox6 and TEML2ox11 (**Supporting Information Fig. S2**). Initially, we examined the timing of growth cessation and bud set by exposing plants to 10 weeks of SD and assessing the bud set scores (**Fig. 2A**). The results indicate that overexpressing *TEML1* and *TEML2*, along with the WT plants, resulted in simultaneous cessation of growth and bud formation (**Fig. 2A**). We examined mRNA levels growth cessation-associated marker genes such as *FT2* and FT2-regulated *GA2ox1* and *GA3ox2* in *TEML1* and *TEML2* overexpressing lines and WT prior to SD photoperiod exposure (Gomez Soto et al., 2022). The result show that none of the evaluated genes exhibited significant differences compared with WT plants confirming that the overexpression of TEMLs genes do not alter mRNA levels and the growth cessation timing (**Fig. 2B and 2C**).

**Figure 2.**
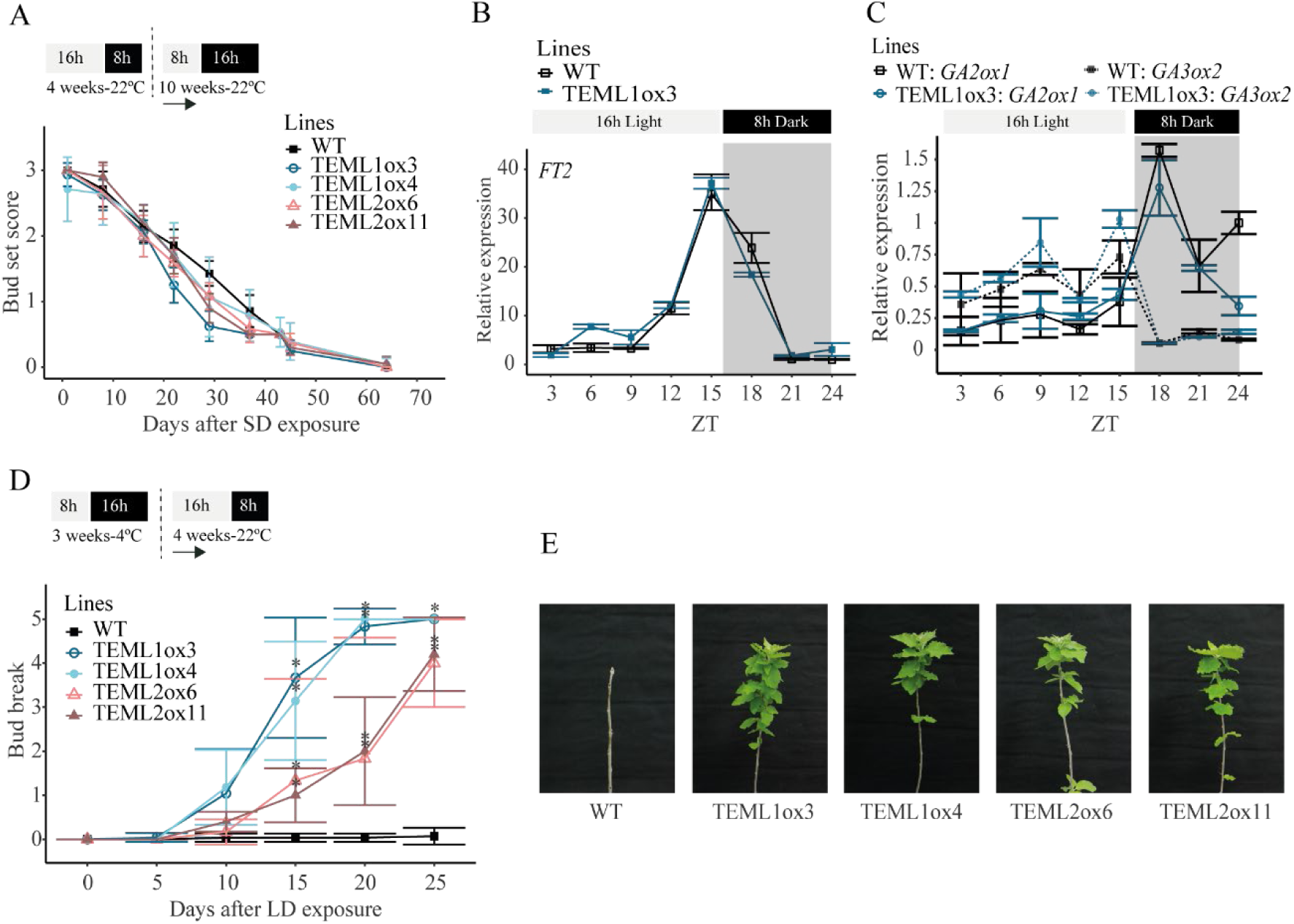
TEML1ox and TEML2ox plants showed early bud break response in endodormant plants. (A) Bud set score of n=15 plants per line measured after subjecting plants to SDs for 70 days. qRT-PCR analysis of relative expression of *FT2* (B) and *GA3ox2* and *GA2ox1*(C) under LD conditions in poplar leaves of TEML1ox and WT plants. UBQ7 was used as housekeeping gene. Plotted values and error bars are fold-change means + s.d. of two biological replicates (D) Bud burst score of TEML1ox and TEML2ox. Values represent the mean score of n = 7 plants after transferring plants from 3 weeks of SD at 4°C to LD at 22°C. (A, D) Significant differences among lines and WT were analysed using Tukey test, *p < 0.05. The top panels indicate photoperiod in which grey boxes mean hours of light and black boxes mean hours of dark, temperature, and time under those conditions. Arrows represent the point at which the experiment began. (E) Representative photos of overexpressing lines taken at the end of the bud break experiment.

In Arabidopsis, the overexpression of *TEM1* or *TEM2* represses the expression of *FT* and delays the transition to flowering by directly binding to the *FT* promoter (Castillejo & Pelaz, 2008; Hu et al., 2021). It has been shown that the overexpression of perennial orthologs of TEMs, *EjRAV1* and *EjRAV2* from loquat (Eriobotrya japonica), also represses *FT* transcription and delays the initiation of Arabidopsis flowering, suggesting a similar function to the Arabidopsis TEMs. However, the overexpression of *EjRAV1* and *EjRAV2* has not yet been tested in loquat (Peng et al., 2021). Arabidopsis TEMs and woody perennials TEMLs proteins exhibit significant similarities in their protein sequences (**Supporting Information Fig. S3**). However, the poplar proteins TEML1 and TEML2 do not retain the ability to repress *FT2* and consequently delay growth cessation and bud set. This difference may be attributed to their lack of interaction with the *FT2* promoter.

In Arabidopsis the AP2 and B3 domains of TEM proteins bind to specific tandem arrange of AP2 and B3 binding elements in the 5’UTR region of the gene, CAACA and CACCTG motifs, respectively (Castillejo & Pelaz, 2008; Hu et al., 2021). The simultaneous association of these two TEMPRANILLO binding domains enhances the transcription factor’s attachment to *FT* promoter (Hu et al., 2021). An *in-silico* analysis of the *FT2 Populus alba* promoter and its 5’ UTR region revealed the absence in this species of the specific tandem arrangement of AP2 and B3 DNA elements found in Arabidopsis *FT* promoter (Castillejo & Pelaz, 2008, **Supporting Information Fig. S4**). However, tandem AP2 and B3 binding sites have been found in the 5′UTR sequence of *Populus tremula* (**Supporting Information Fig. S4D)**. This lack of *FT2* repression in our poplar TEML overexpressing lines may be due to the inability of TEMLs to bind the *FT2 Populus alba* promoter with high affinity. Consequently, the transcriptional regulation of hybrid poplar *FT2* promoter appears to have lost its tandem arrange of TEMPRANILLO regulatory elements and associated repressive function.

### *TEML1* and *TEML2* overexpression breaks poplar endodormancy

We then investigated whether *TEML1* and *TEML2* are involved in regulating the endodormancy stage. After the growth cessation and the bud set, all TEMLox lines together with WT plants were subjected to an insufficient chilling treatment of 3 weeks in SD conditions at 4°C. Afterward, they were transferred to LD conditions at 22°C to assess bud break. While WT plants neither broke buds nor resumed growth indicating they remained in endodormancy, *TEML1* and *TEML2* overexpressing lines displayed bud break, with bud initiation occurring after 10 days of LD (**Fig. 2D and 2E**). The results indicate that lines overexpressing *TEML1* and *TEML2* are unable to maintain the endodormancy stage. This inability could be due to the transgenic plants not reaching this stage deeply enough or because the overexpression of TEML affects the duration of endodormancy. These findings suggest that *TEML1* and *TEML2* may play a role in maintaining endodormancy, and their upregulation could lead to an early release from this phase.

### The overexpression of *TEML1* and *TEML2* primarily affects the transcriptome associated with endodormancy

To understand the molecular changes in *TEML1* and *TEML2* overexpression lines during endodormancy, we conducted a comparative RNAseq analysis on the apical buds of *TEML1* overexpressing, *TEML2* overexpressing, and WT plants at two different stages: bud set and endodormancy. To achieve this, we formed two distinct groups: plants in the bud set stage that were treated with 8 weeks of short days (SD8W point), and plants in endodormancy stage that received 10 weeks of short days followed by 2 weeks of chilling (CH2W point). Transcriptomes were analysed to obtain lists of differentially expressed genes (DEGs) by comparing each *TEML* overexpressing line with its WT control and then combining the results to identify common response genes for each time point. Our analysis identified 35 common DEGs at the SD8W and 444 common DEGs at the CH2W (**Fig. 3A and 3B**). When comparing these two lists, we identify only 3 overlapping DEGs, with 32 DEGs expressed exclusively at the SD8W time point and 441 DEGs specific to the CH2W time point (Figure 3C). The minimal molecular differences in SD8W indicate that lines overexpressing *TEML1* and *TEML2* enter dormancy in a similar manner to the wild type, whereas the bigger transcriptomic changes observed in CH2W indicate a failure during the endodormancy period in *TEML* overexpressing lines.

**Figure 3.**
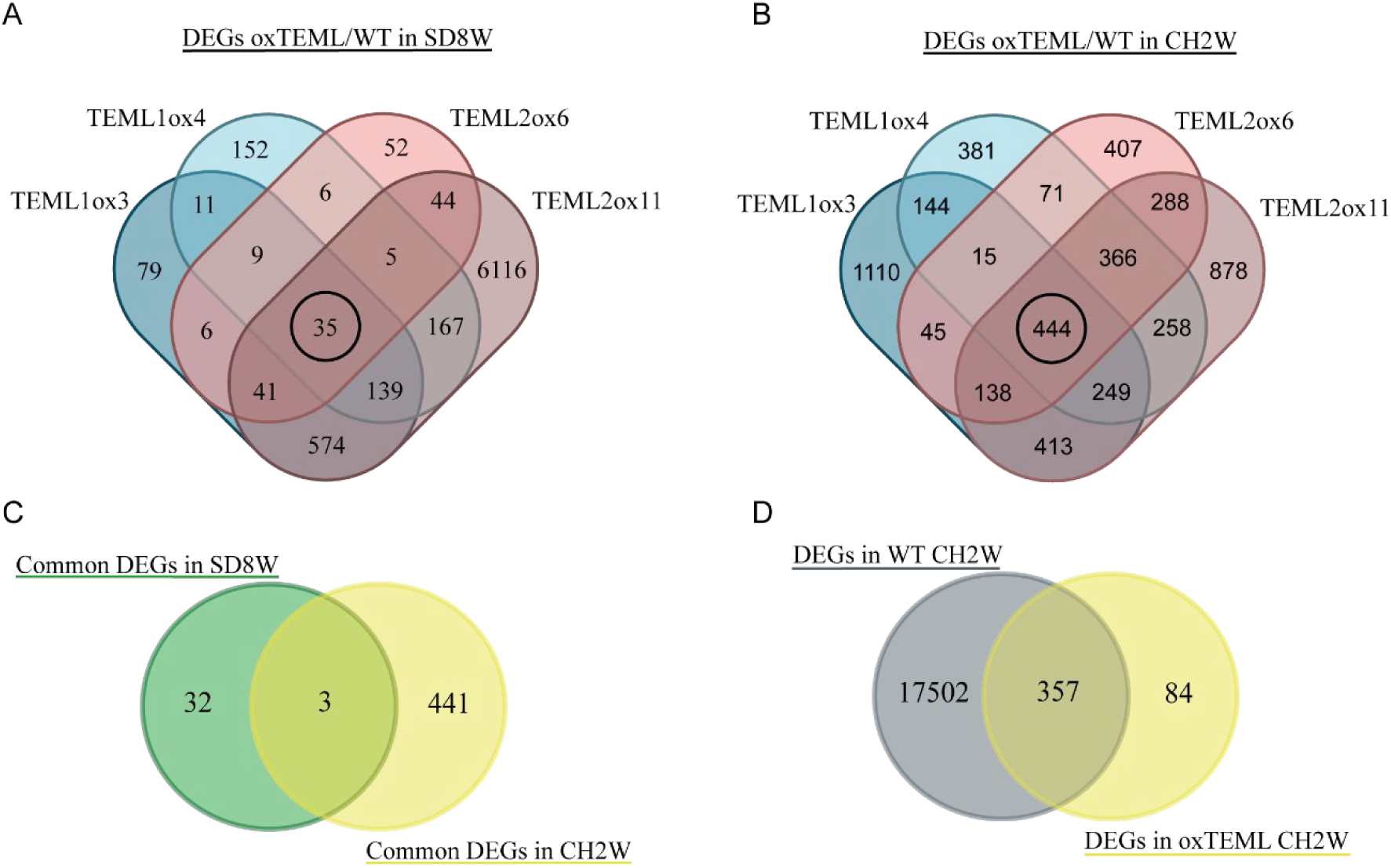
Venn diagrams representing common DEGs of TEML1ox and TEML2ox buds. Apical buds from TEML1 and TEML2 overexpressing lines and WT plants were collected at zt7 after 8 weeks of SD exposure (SD8W) and 8 weeks of SD exposure followed by 2 weeks of chilling (CH2W). DEG lists were obtained by comparing TEML overexpression lines to their respective WT controls under SD8W (A) and CH2W (B) conditions using RNAseq analysis. Two replicates were analysed per genotype. The inner circle in the Venn diagrams represents the common DEGs of all the overexpressing lines (A,B). The Venn diagram in panel (C) overlaps common SD8W DEGs with common CH2W DEGs (C). Panel (D) shows a Venn diagram comparing specific CH2 genes, obtained by comparing DEGs in WT buds under CH2W and DEGs in WT buds in SD8W with the 441 common DEGs in CH2W from the overexpressing plants (D). RNAseq data comparison and DEG lists were generated using the EdgeR package with a threshold of *p < 0.05.

Afterwards, we identified specific chilling-related genes in WT plants by comparing the DEG list from CH2W to SD8W. We identified 17502 genes that significantly changed their expression after chilling exposure, 48% of them were upregulated and 52% were downregulated (**Supplementary Table 2**), showing the huge transcriptional rearrangement that occurs when buds become endodormants. We then combined this list of chilling responsive genes with the 441 common DEGs identified in the *TEML* overexpressing lines at the CH2W time point (**Fig. 3D**). Among these, we found an overlapping of 357 DEGs, indicating their specific role during the endodormancy (**Fig. 3D**). The expression of chilling-specific 357 DEGs displayed opposite patterns compared to WT plants, clustering in two 2 main groups: one comprising upregulated genes and the other consisting of downregulated genes in the overexpressing lines compared with WT (**Supporting Information Fig. S5**). These genes are both altered during dormancy and associated with the early bud break phenotype in *TEML1* and *TEML2* overexpressing plants.

### *TEML* overexpressing lines cause an early induction of growth reactivation genes

To identify the functional relationships and biological relevance of the chilling-specific 357 DEGs identified, we conducted GO terms network analysis. We separated the datasets into upregulated and downregulated genes, performing GO enrichment analysis on each group independently. The analysis revealed that the 212 upregulated genes were associated with 79 GO terms, while the 131 downregulated genes were linked to only 10 GO terms (**Supporting Information Fig. S6**). Our GO network analysis produced four distinct interconnected subgroups within the upregulated dataset and only one network in the downregulated dataset (**Fig. 4 A, C, E, G, I**). Among the upregulated genes, the first subgroup focused on processes related to chromatin remodeling and organization, heterochromatin formation, DNA methylation and regulation of gene expression (**Fig. 4A**). The second subgroup is linked to the activation of shoot development (**Fig. 4C**). The third subgroup encompasses primary metabolic processes related to cell growth reactivation such as macromolecule metabolism, macromolecule modification, protein phosphorylation and nitrogen compound metabolism (**Fig. 4E**). Finally, the fourth subgroup includes general GO categories and illustrates the connection between developmental and metabolic processes (**Fig. 4G**). Following, we examined the temporal expression patterns of all genes within each network using our previously published transcriptomic dataset from poplar apical buds of plants grown under natural conditions, from “Mid Winter f1” to “Mid Spring” (Conde et al., 2019). More than 90% of the genes in each subgroup showed specific inducible expression from mid winter to mid spring, with 50% of those genes being specifically activated in the spring (**Fig. 4B, D, F, H**). Collectively these analyses indicate that overexpression of TEML activates genes and processes related to growth reactivation after two weeks of chilling exposure, which could be required for the early bud break phenotype observed.

**Figure 4.**
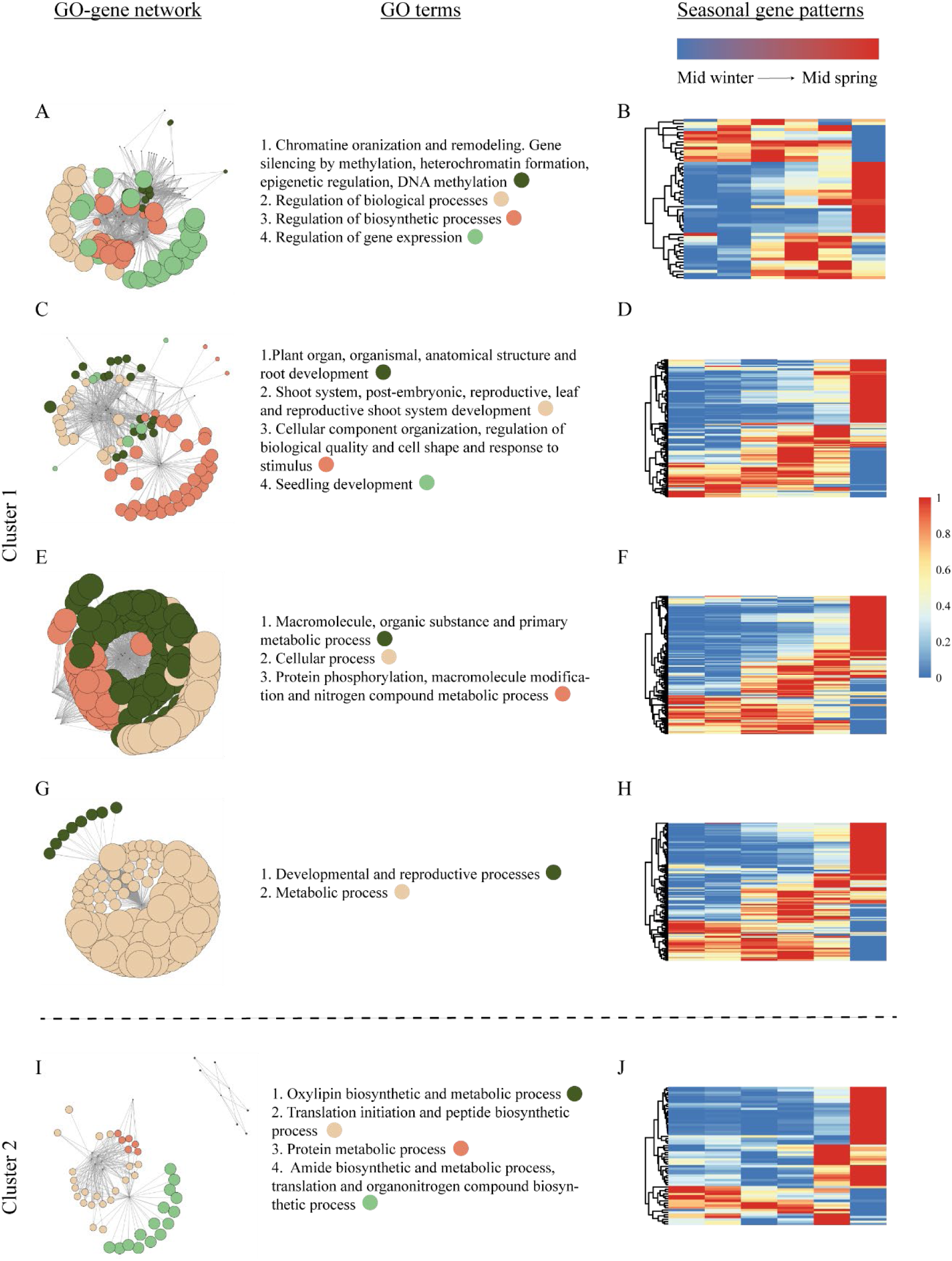
GO enrichment analysis of the 357 DEGs during endodormancy. Networks (left panels) represent upregulated (A-G) and downregulated (I) GO terms and genes during CH2W. Using Arabidopsis annotation, we obtained the relationships between GO terms and GO-to-gene to build the network. We clustered the network by applying the Louvain method for community detection, and we assigned unique colors to clusters (including GO terms and genes).The size of each node reflects the number of genes linked to a given GO term, with larger nodes representing a higher number of associated genes, indicating a directly proportional relationship. Each subgroup was clustered based on the internal network relationships and since all genes and GO terms were interconnected, they formed a single unified network. (B, D, F, H, J). The middle panels describe the most characteristic GO term group in each network. Heatmaps (right panels) illustrate the natural expression of DEGs in each subgroup. Values are normalized data of apical bud RNA collected from Mid Winter to Mid Spring, “Early”, “Mid” and “Late” indicate the first, second and third month of each season, and “f1” and “f2” indicate the fortnight 1 or 2 of the month. The colour panel above the heatmaps represents the highest expression levels in red and the lowest levels in blue.

A similar analysis was conducted for the downregulated genes and their corresponding enriched GO terms. The enrichment network analysis identified a group associated with secondary metabolic processes that were not linked to the earlier results. Additionally, the network connects genes and GO terms related to protein translation and the biosynthesis of organonitrogen compound (**Fig. 4I**). The genes within this network also exhibit a temporal profiling displaying specific inducible expression from mid winter to mid spring in the apical buds of poplar. The downregulation of those genes in TEML overexpressing lines, following two weeks of chilling treatment, diminishes their implication in early endodormancy exit phenotype observed.

### Chilling-induced *TEML1* and *TEML2* promote endodormancy exit

To gain a better understanding of the molecular functions that reduce the dormant stage in lines overexpressing *TEML1* and *TEML2*, we examined the presence of key known regulators of endodormancy among the 212 upregulated genes. Among them, we identified several genes that may contribute to shortening dormancy by reducing the chilling requirement (**Fig. 5A**). We found two poplar orthologs of *BRASSINOSTEROID INSENSITIVE 1 (BRI1)*, whose overexpression in Arabidopsis eliminates the need for cold stratification in seed germination (Kim et al., 2019). Additionally, we identified a poplar ortholog of *VERNALIZATION INSENSITIVE 3 (VIN3)*. Upregulating this gene in Arabidopsis can shorten the vernalization period, depending on the duration of exposure to cold temperatures (Hepworth et al., 2018). The presence of poplar homologs to the Arabidopsis regulators of vernalization shows how evolutionarily related regulatory mechanisms may be adopted for different processes such as seed and bud dormancy, both requiring a certain period of chilling accumulation to release dormancy (Yang et al., 2021). Furthermore, we discovered a poplar ortholog of *GROWTH-REGULATING FACTOR 7 (GRF7)*, which, upon induction, represses ABA signalling to prevent growth inhibition under osmotic stress conditions (Kim et al., 2012). Proper regulation of ABA metabolism and signaling is necessary to maintain the dormancy. Poplar SVL induces ABA biosynthetic genes and also acts downstream of the ABA pathway, forming a positive feedback regulation with ABA signalling and mediating dormancy establishment and maintenance (Singh et al., 2018, 2019; Tylewicz et al., 2018). Thus, GRF7 activation might be necessary to promote the growth resumption by repressing ABA signaling in poplar buds.

**Figure 5.**
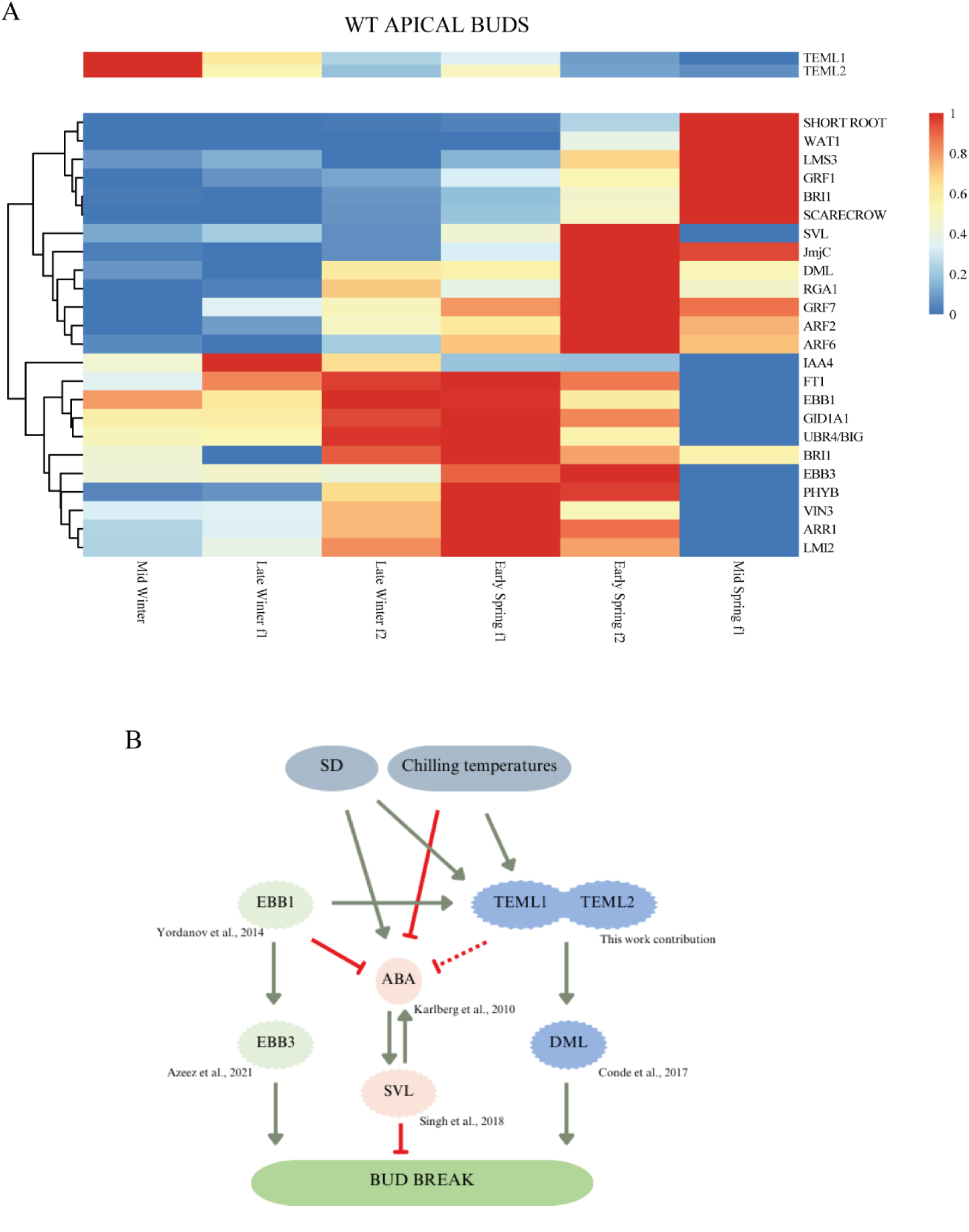
Proposed model of TEML1 and TEML2 roles during chilling. (A) Seasonal expression patterns of known regulators of endodormancy and growth in apical buds. *TEML1* and *TEML2* are shown in the upper panel. Gene candidates proposed to shorten dormancy period, including *FT1*, *EBB1* and *EBB2* are shown in the bottom heatmap. Values represent normalized RNA data from apical buds collected from Mid Winter to Mid Spring, “Early”, “Mid” and “Late” indicate the first, second and third month of each season, and “f1” and “f2” indicate the fortnight 1 or 2 of the month. (B) Proposed model of bud break regulation. Environmental conditions are shown in gray. EBB1 pathway is shown in green, while ABA-SVL pathway is shown in pink. The proposed TEML1 and TEML2 pathway, along with its connections to the described pathways, is depicted.

Additionally, we identified a set of genes whose induction is necessary to promote shoot apex growth. PHYTOCHROME B (PHYB) and DEMETER (DEM) have already been shown to regulate poplar bud break (Conde et al., 2017). The gibberellin signaling genes *GID1A* and *RGA1* have been associated with the exit from dormancy in perennials (Yue et al., 2018). Furthermore, a poplar ortholog of the Type B cytokinin response regulator ARR1 is required for shoot meristem activity in Arabidopsis (Liu et al., 2020). Moreover, we discovered a group of auxin response genes, including *ARF2*, *ARF6*, *IAA4*, *SHORTROOT*, *SCARECROW*, *BIG*, and *WAT1*, that may be essential for creating patterns for organ differentiation from the shoot meristem niche (Catalá et al., 2019; Conde et al., 2017; Ding et al., 2021; Hepworth et al., 2018; Howe et al., 2015; J.-S. Kim et al., 2012; S. Y. Kim et al., 2019; Pastore et al., 2011; Ranocha et al., 2013; Salvi et al., n.d.; Shimano et al., 2018; Yue et al., 2018). Recently, activation of auxin response has been associated to bud break through temperature-mediated chromatin remodelling during winter dormancy in apple (Chen et al., 2022). Apart from phytohormone regulators, we found activation of Jumonji-domain-containing transcription factor JUMONJI 27 that shows H3K9 histone demethylase activity and regulation of flowering time in Arabidopsis (Dutta et al., 2017). We also found a poplar ortholog of the LATE MERISTEM IDENTITY2 (LMI2) activator APETALA1, which promotes shoot organogenesis in Arabidopsis (Pastore et al., 2011). Overall, the increased expression of these genes in TEML1 and TEML2 overexpressing plants at CH2W suggests a shoot meristem that is overcoming a dormant state.

Our study indicates that *TEML1* and *TEML2* are transcription factors induced by short days (SD) and chilling temperatures (**Fig. 1C and 1F**). During the seasonal transition from “Mid Winter” to “Mid Spring,” we observed that the expression levels of *TEML1* and *TEML2* are highest in Midwinter, gradually decreasing as we move toward spring (**Fig. 5A**). The expression peaks of the TEML-regulated genes *BRI*, *VIN3*, and *GRF7* follow the peaks of *TEML1* and *TEML2* (**Fig. 5A**) indicating their activation could be dependent on *TEML*. Additionally, the expression peak of TEML-regulated genes associated with shoot meristem reactivation occurs after the mid-winter time point (**Fig. 5A**). Overall, these findings support the role of *TEML1* and *TEML2* in regulating the activation of pathways that could contribute to chilling fulfilment and the reactivation of growth. Our data suggest that *TEML1* and *TEML2* act upstream of the previously reported *DEMETER* pathway in poplar (Conde et al., 2017). The mining of the *EBB*1 datasets shows that TEML1 and TEML2 are upregulated in plants overexpressing EBB1 (Yordanov et al., 2014), suggesting that EBB1 could act in concert with TEML1 and TEML2 to promote poplar bud break.

## CONCLUSION

TEMPRANILLO transcriptional function might undergo neofunctionalization in hybrid poplar. Lines that overexpress the poplar genes *TEML1* and *TEML2* do not suppress *FT2* transcription or delay the timing of seasonal growth cessation and bud set. *TEML1* and *TEML2* are induced under winter conditions, specifically during short days and chilling temperatures. Furthermore, lines that overexpress *TEML1* and *TEML2* specifically alter the endodormancy transcriptome and facilitate the breaking of endodormancy. We propose that short days and chilling conditions activate *TEML1* and *TEML2*, which in turn activate genes that limit the duration of endodormancy and promote vegetative development of the shoot apical meristem (**Fig. 5B**).

## Acknowledgements

The study was supported by grants PID2021-123060OB-I00 awarded to MP and IA and the Severo Ochoa (SO) Programme for Centres of Excellence in R&D from the Agencia Estatal de Investigación of Spain [grant CEX2020-453 000999-S (2022 to 2025) to the CBGP]. DGS has a FPI fellowship (PRE2019-089312) from Ministerio de Ciencia e Innovacion de España. PMT is supported by Scuola Superiore Sant’Anna. DC has a Caixa Junior Leader Fellowship.

## SUPPLEMENTAL FIGURES

**Supplementary Figure S1.**
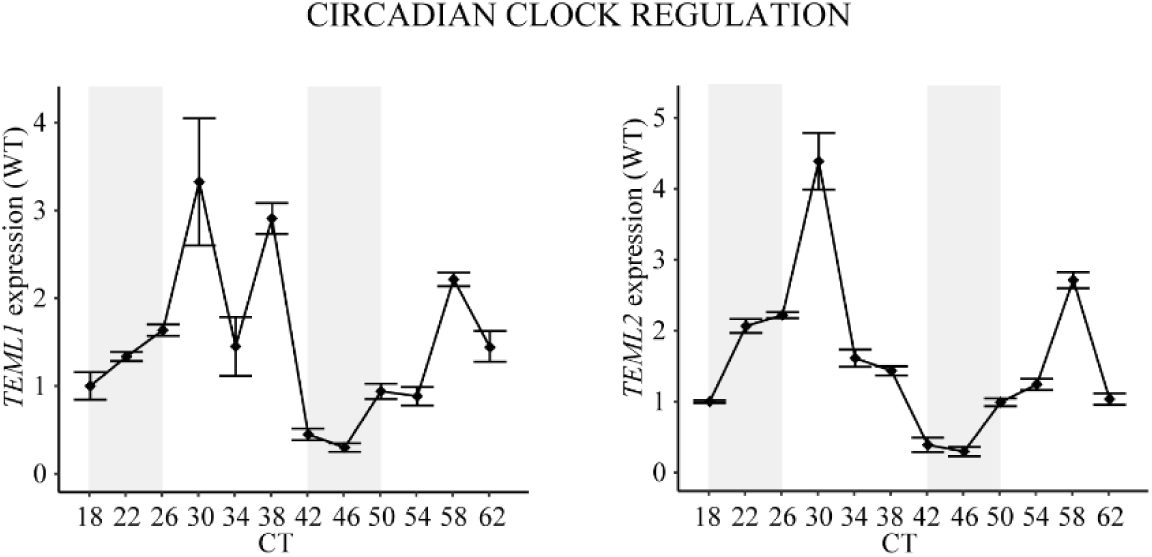
Relative expression of *TEML1* (A) and *TEML2* (B) under constant light (LL) in WT plants. Grey panels denote the time corresponding to the nighttime phase of the experiment. The x-axis means the circadian time (CT). *UBQ7* was used as housekeeping gene. Plotted values and error bars are fold-change means + s.d. of two biological replicates.

**Supplementary Figure S2.**
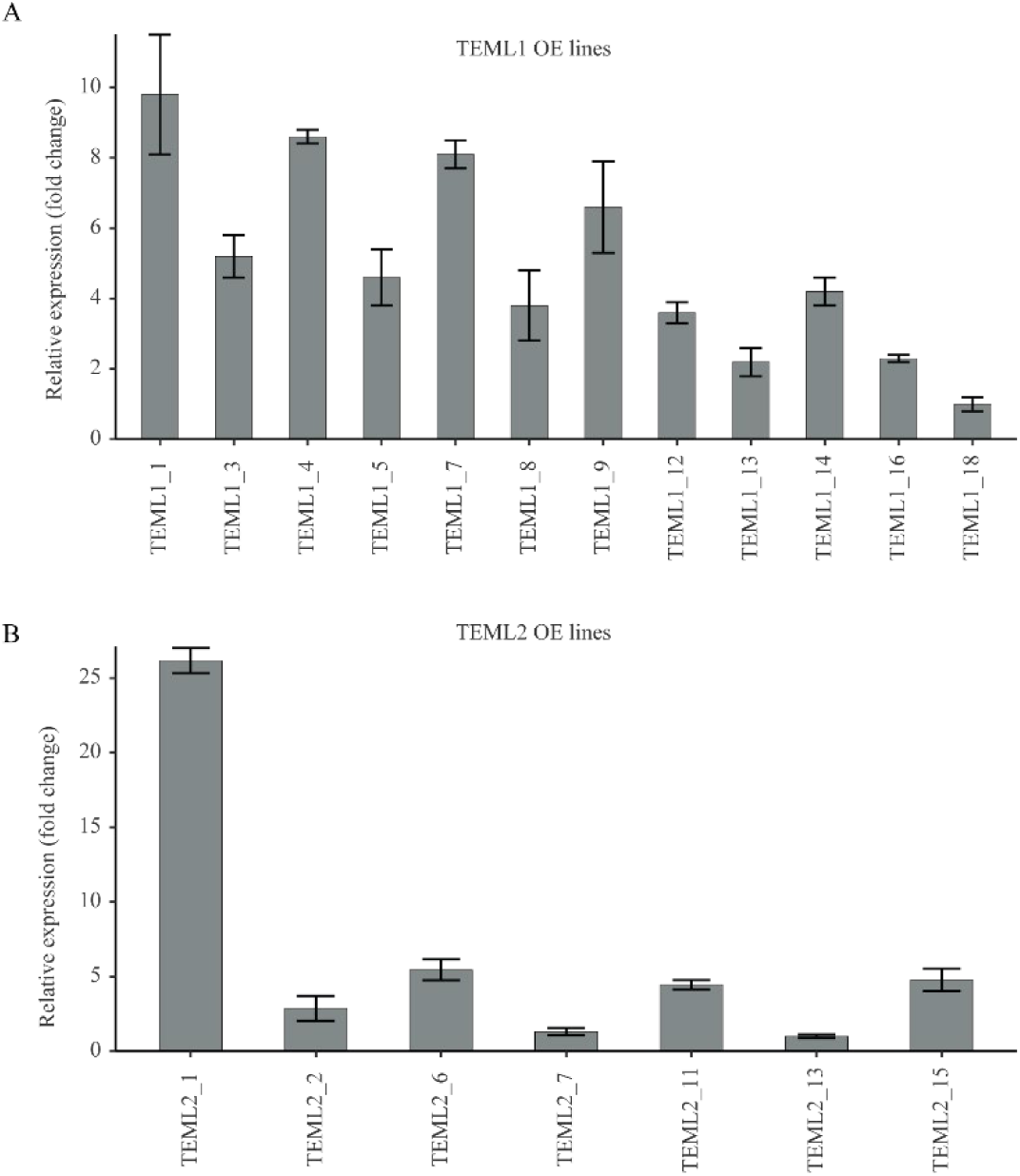
Characterization of *TEML1* and *TEML2* transgenic lines. qRT-PCR analysis of *TEML1* and *TEML2* overexpressing plants. *Ubiquitin7* is used as the housekeeping gene. Plotted values and error bars are fold-change means ± s.d. of two biological replicates.

**Supplementary Figure S3.**
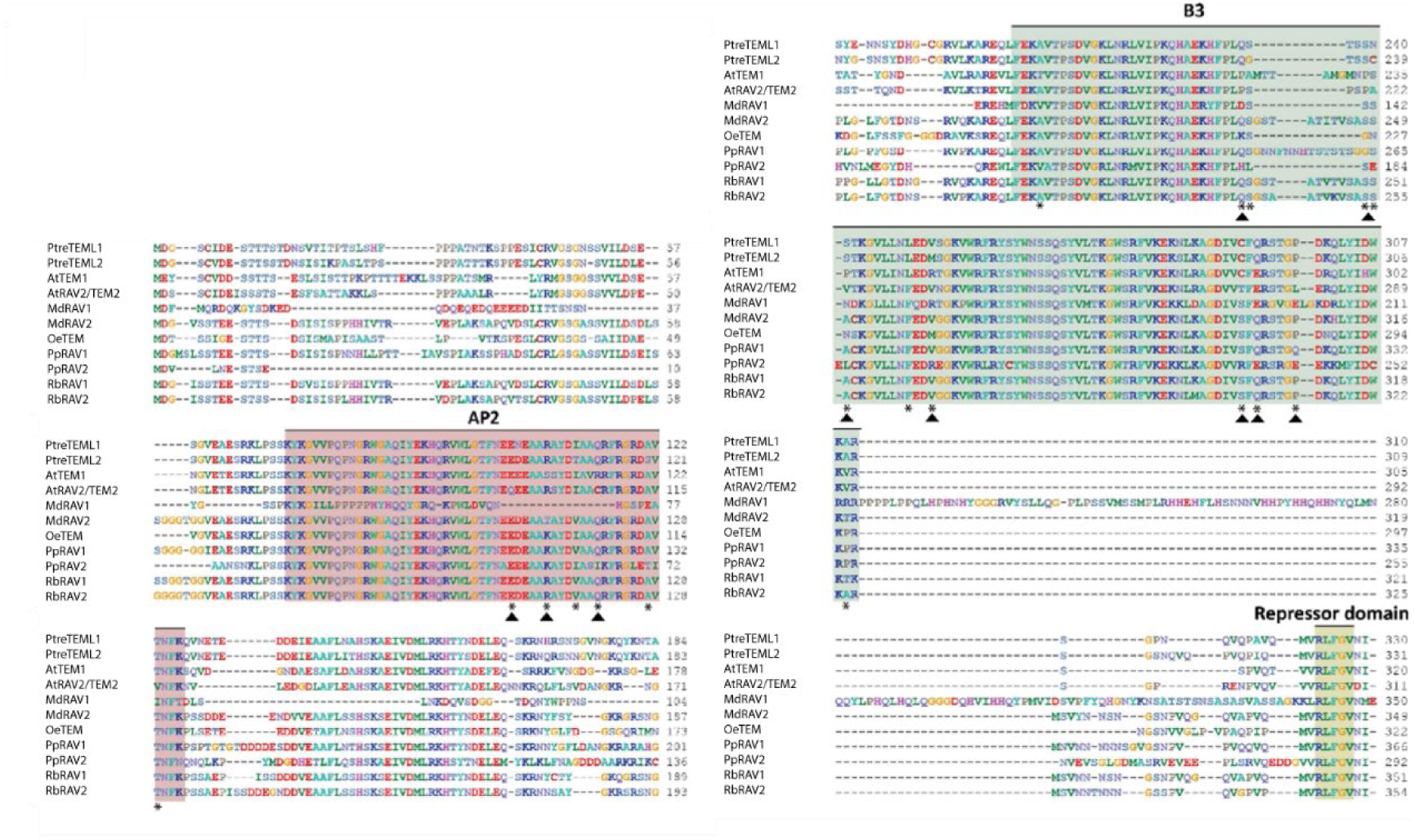
Clustal X alignment of RAV and TEM peptide sequences in poplar (Ptre), Arabidopsis (At) apple tree (Md), olive (Oe), peach tree (Pp) and loquat (Rb). Characteristic family domains: AP2, B3 and repressor domain are highlighted with pink, green and yellow background, respectively. Asterisks (*) indicate amino acid variations in the sequence. Black arrows point to physiologically significant amino acid changes.

**Supplementary Figure S4.**
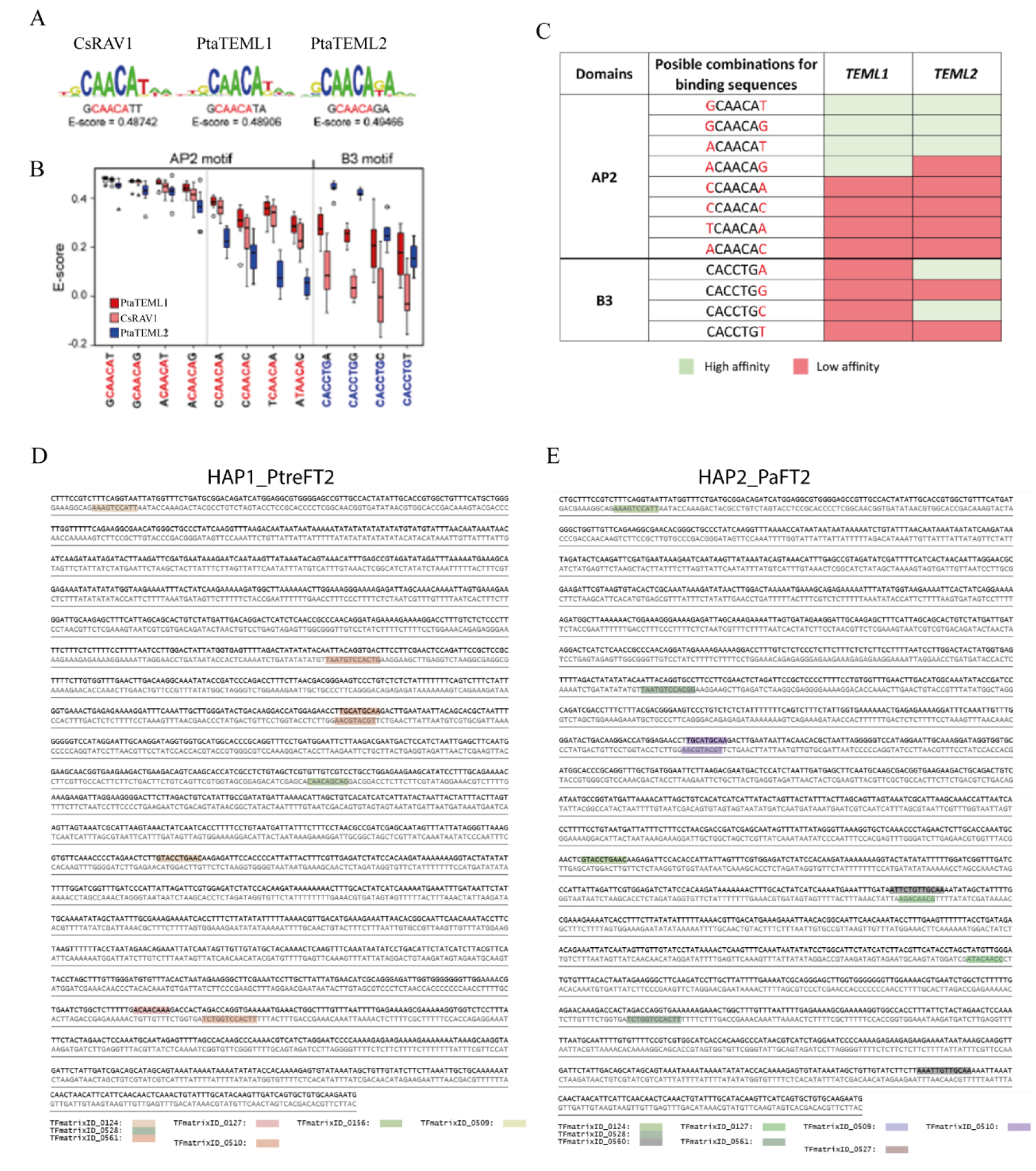
TEML binding sequences obtained by Protein Binding Matrix and *FT2* promoter and 5’ UTR AP2 and B3 elements. (A). Position weight matrix (PWM) representation of the top scoring 8-mers obtained for chestnut CsRAV1, and poplar PtaTEML1 and PtaTEML2 proteins (B) Box-plot of E-scores of the AP2 and B3 DNA elements Boxes in blue correspond to full length PtaRAV2 (PtreTEML2), red boxes represent PtaRAV1 (PtreTEML1), Orange ones correspond to CsRAV1 proteins expressed in E. coli. (C) Summary table containing PtreTEML1 and PtreTEML2 AP2 and B3 DNA binding domains and their different combinations of flanking base pairs. Third and fourth columns represent their binding affinity to said domains. Colour code represents high (green) or low affinity (red). (D) FT2 promoter alignment of *Populus tremula* (HAP1) highlighting the AP2 and B3 boxes. (E) FT2 promoter alignment of *Populus alba* (HAP2) similarly highlights the AP2 and B3 boxes.

**Supplementary Figure S5.**
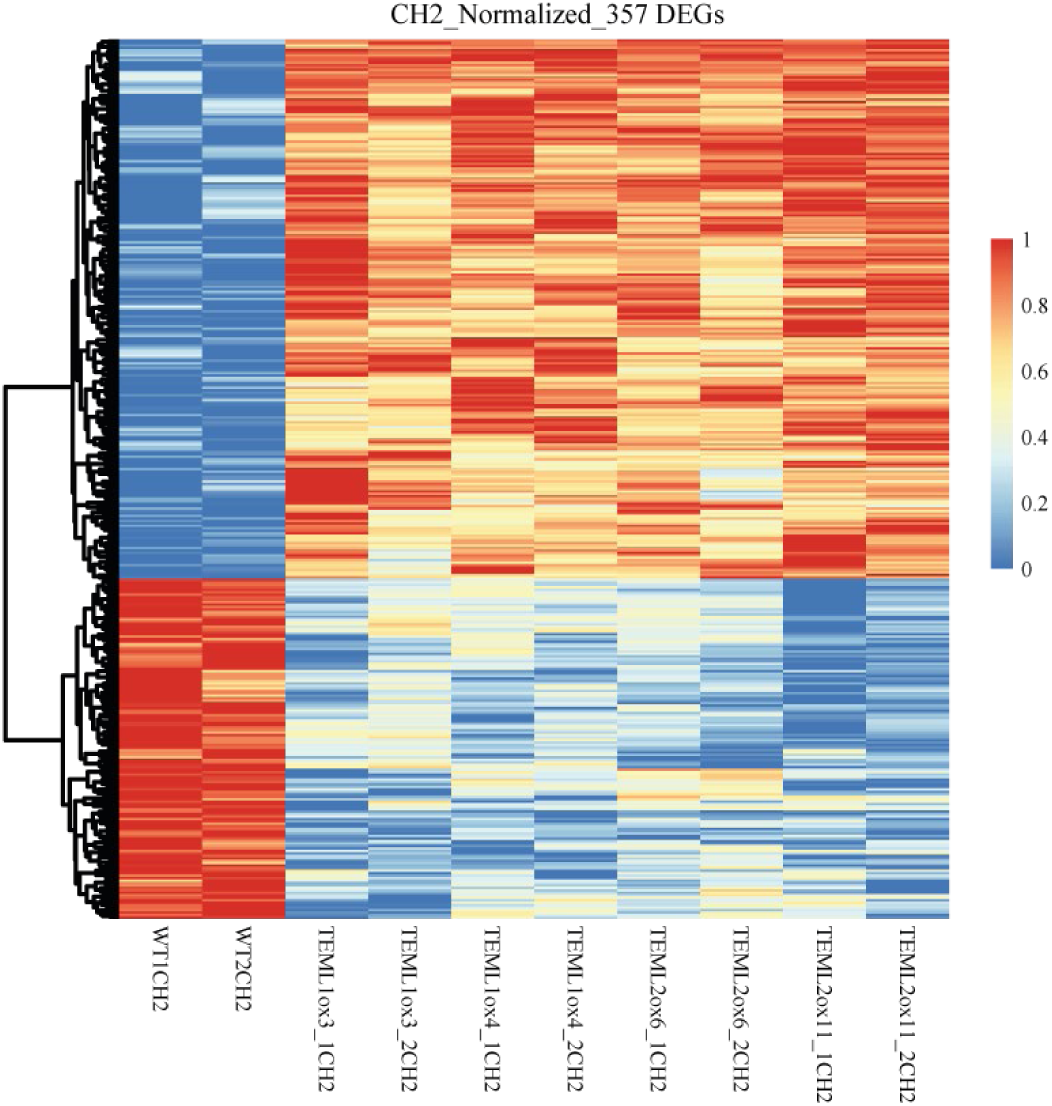
Heatmap illustrates normalized data of the 357 chilling specific common DEGs. Bottom labels indicate the transgenic line shown in the upper column. Two biological samples of each genotype were analysed.

**Supplementary Figure S6.**
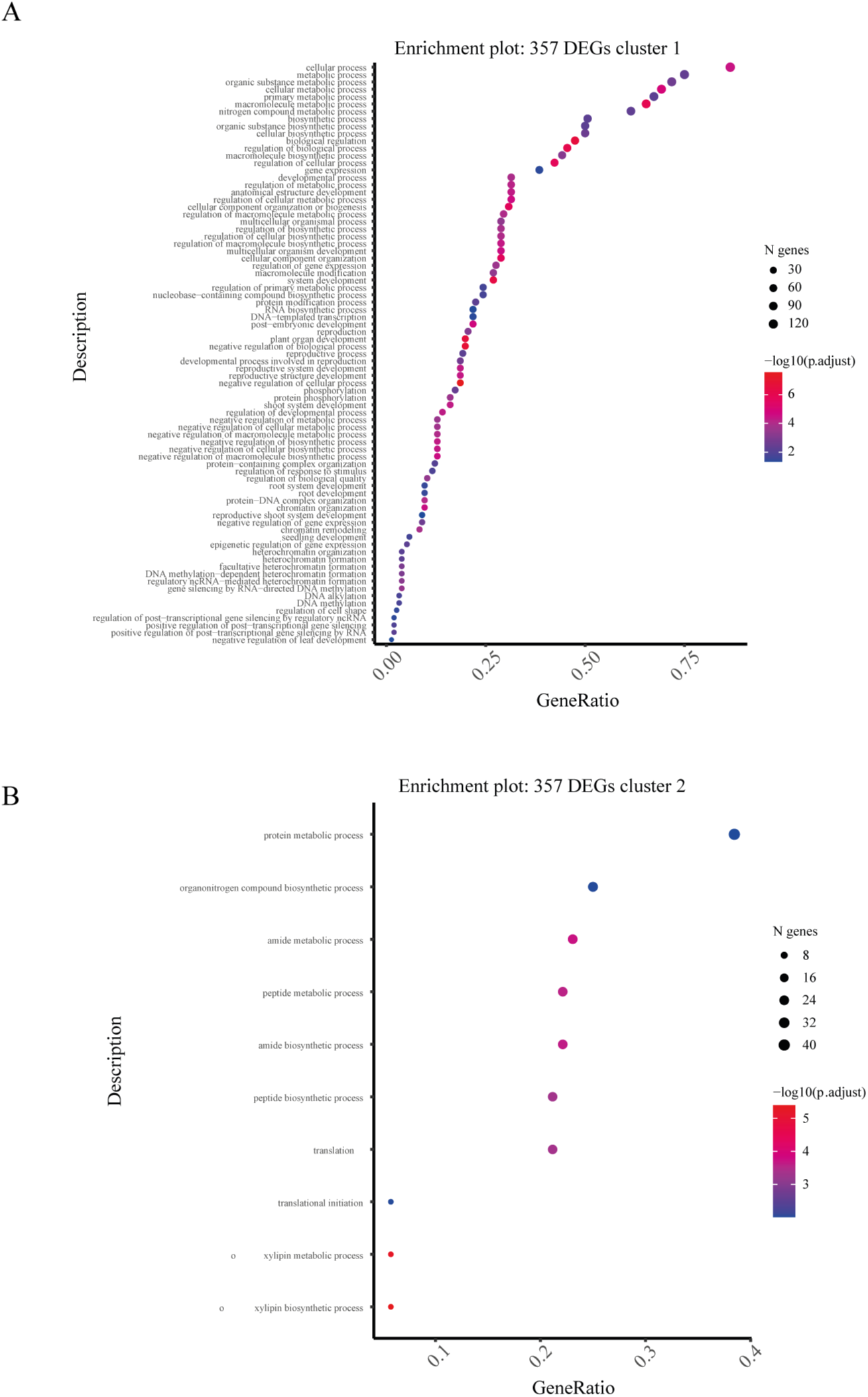
GO terms associated with upregulated and downregulated genes. Analysis was performed using gProfiler tool and selecting only Biological process for this study. P value < 0.05.

